# Creation of Novel Methylating Conjugal Donor Strains for Genetically Recalcitrant Bacteria

**DOI:** 10.64898/2026.01.22.701203

**Authors:** A W Dempster, R Mansfield, P Ingle, Z Bean, C Akaluka, J Millard, N P Minton

**Affiliations:** Synthetic Biology Research Centre, University of Nottingham, UK

## Abstract

A methodology is presented to integrate DNA methyltransferase genes of interest into the single copy, IncPβ ‘R’ Factor conjugal plasmid, R702. This is to enable DNA transfer, or improved frequency thereof, by mirroring the recipient native DNA methylation signature prior to transfer of plasmid DNA by conjugation, evading native restriction modification systems in the organism of study, without target strain modification. As proof of concept, the bacteriophage derived methyltransferase Ф3TI, known to facilitate DNA transfer into the industrially important model solventogenic organism, *Clostridium acetobutylicum* ATCC 824, was inserted into the single copy R702 to create an *in vivo* methylating strain. Plasmids extracted from this strain were transferred by electroporation at a comparable frequency to the established method which uses the multi-copy accessory plasmids pAN1 or pAN2, harbouring Ф3TI. The methodology was further exemplified and employed to enable high frequency of DNA transfer into the clinically relevant *Clostridioides difficile* ribotype 027 outbreak strain, R20291. The modification (*hsdM*) and specificity (*hsdS*) components of the native Type I restriction modification system of R20291 were inserted into R702 to create a functional methylating conjugal donor *Escherichia coli* strain. This approach of coupling methylation and the native transfer functions of R702 creates a system which can easily be mobilised into a genotypically appropriate *E. coli* strain; creating an *in vivo* methylating donor strain which may be utilized to protect and transfer DNA to the organism of study by conjugation.

## 1. Introduction

Since Frederick Griffith’s first observations of bacterial transformation in 1928, and the subsequent elucidation and confirmation of DNA as Griffith’s “transforming principle” in 1944 by Avery and colleagues and Hershey and Chase in 1952; transformation has been a tool critical to the advancement of molecular biology^1,2,3^. Griffith’s seminal experiments are an example of natural transformation, the active uptake of exogenous DNA by a bacterium from its environment. In nature, horizontal gene transfer may also occur by bacteriophage transduction, or the cell to cell transfer of DNA by bacterial conjugation^4^. Conjugation, the most probable horizontal gene transfer event in nature, requires temporal and spatial stringencies of viable donor and recipient cells, conditions which can be replicated in the laboratory to introduce DNA into an organism of study. Donor cells must possess a fertility factor, ‘F’, which encodes the functions necessary for conjugative DNA transfer, the *tra* genes^5^. The F factor is often contained on a replicative and self-mobilising episome, which may also be chromosomally integrated^6^.

The ‘F’ plasmids are the established paradigm for conjugative DNA transfer in Gram-negative bacteria, with the closely related ‘R’ factor plasmids associated with carriage of resistance factors, as well as encoding transfer functions. The IncPβ R factor plasmid, R702, is routinely used as an accessory plasmid for conjugal DNA transfer from *E. coli* into *Clostridium* and other bacterial genera ^7, 8^. Isolated from a strain of *Proteus mirabilis*^9^, the published DNA sequence indicates a large plasmid of 76,587 base pairs in size, harbouring genes encoding resistance mechanisms for spectinomycin, tetracycline, kanamycin, sulphonamides and mercury salts, as well as the *tra* operon, necessary for conjugal gene transfer^10^.

Horizontal incursion of non-host, parasitic DNA may present a threat to cell viability through the expression of lethal genes or genomic aberration, and so DNA uptake and transfer may be constricted by protective host factors, impeding molecular methods in the organism of study. Host defence mechanisms are diverse, clustering in so called “defence islands” in the microbial genome^11^. The most abundant and obstructive of these defence factors are the restriction modification (RM) systems^12^. These ubiquitous, innate intracellular defence mechanisms recognise and degrade foreign nucleic acid molecules to preserve cellular integrity, whilst host DNA remains intact and potentially beneficial DNA, from related species, is permitted and maintained. To date, four types of RM systems have been described, with distinct mechanisms of recognition and degradation^13^. Broadly, Types I-III are comprised of genes encoding a methyltransferase (MTase) and restriction endonuclease (REase), which cleave DNA designated as ‘non-self’ due to lacking host methylation signatures at specific recognition sequences. The Type IV systems encode methylation dependent REases, cleaving DNA with a non-host methylation pattern^14,15^. Genetically recalcitrant bacteria may harbour one particularly discriminatory system or multiple, presenting significant barriers to gene transfer, impeding the study of these organisms^16^.

Efforts, heretofore, to improve DNA transfer frequency by RM evasion include recipient strain modifications^17,18^ and heat treatment^19^, plasmid DNA site avoidance^20^, *in vivo* methylation prior to transfer^21,22,23^ and the methylation status of the *E. coli* donor strain^24^. Here a methodology is presented to create methylating conjugal donor strains; coupling the activities of DNA methylation and mobilisation, by lambda Red mediated integration of DNA modifying MTase genes into the ‘R’ factor plasmid, R702^25^. The modified R702 plasmid can subsequently be transferred from the recombinant strain into a suitable *E. coli* strain by conjugation or electroporation, to be used as a conjugal donor host. Shuttle plasmids are able to be transferred and maintained in the intended recipient organism, without modification.

## 2. Results

### The Recombineering Base Vector

The recombineering base vector (pRBV1) is designed to harbour MTase gene(s) of interest within a recombineering cassette comprising an AraC-P_BAD_ arabinose inducible promoter, the inserted MTase gene(s) of interest and the aminoglycoside acetyltransferase, *aac*, apramycin selection marker gene. Completing the recombineering cassette are two flanking, 500 bp regions of sequence homology to the *aadA1* gene locus within the R702 plasmid. MTase genes of interest are inserted into the plasmid following linearisation of pRBV1 by PCR or restriction digest. The plasmid backbone is comprised of the erythromycin resistance marker, *ermB*, and Gram-negative replicon, ColEI, from the pMTL80000 vector series^26^. The entire cassette may be amplified with PCR using flanking primers to generate linear DNA for subsequent recombineering steps (Figure 1).

**Figure 1:**
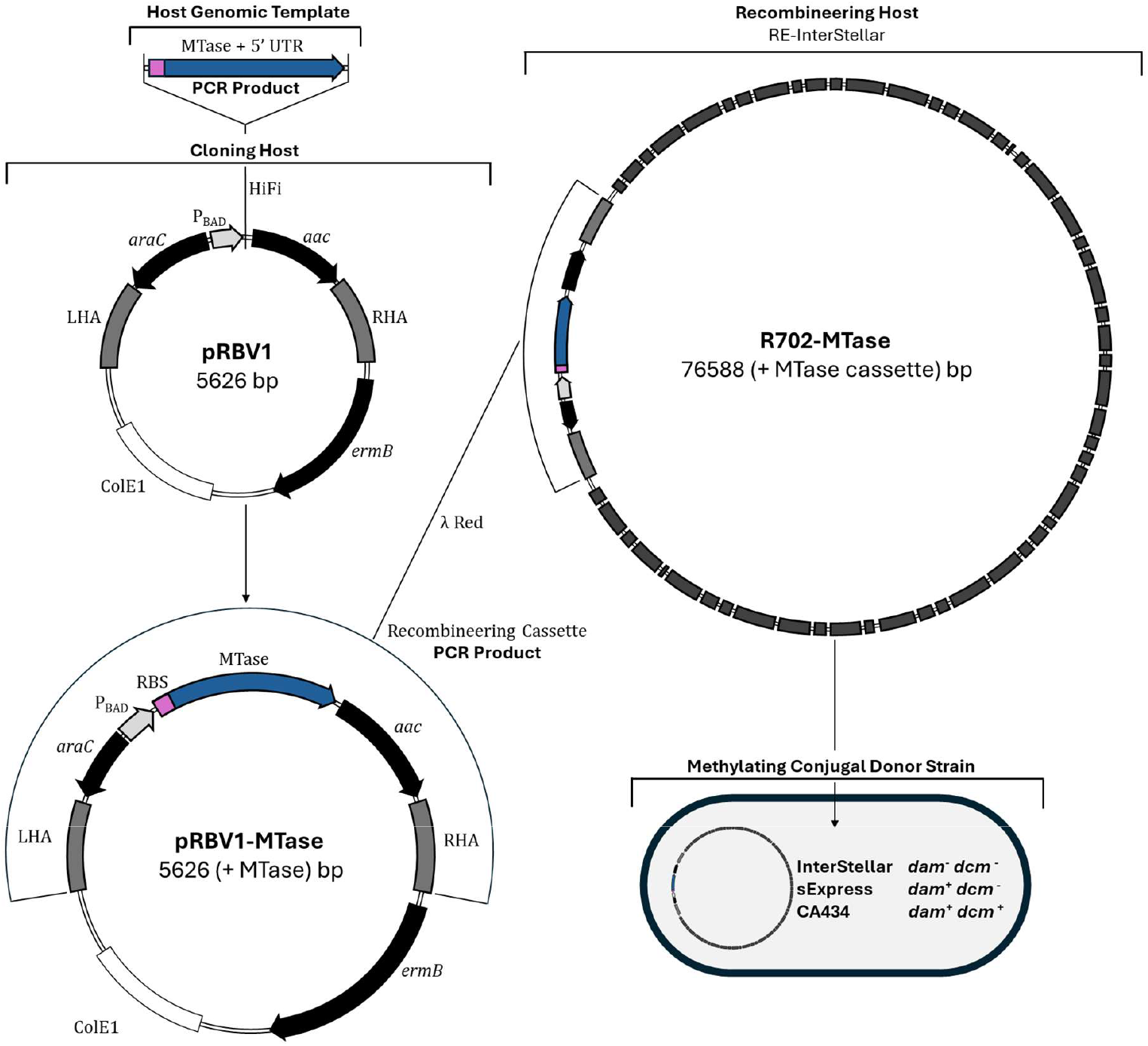
Methylating Conjugal Donor Strain Workflow. The MTase gene of interest is amplified from the RM harbouring strain of study and inserted into the recombineering base vector, pRBV1, by HiFi assembly (NEB) or restriction cloning. The entire recombineering cassette containing an arabinose inducible promoter, MTase and apramycin resistance marker, is PCR amplified along with flanking homology arms, purified on agarose gel, extracted and transformed into the RE-InterStellar recombineering host. Recombinant clones are selected for by growth on plates supplemented with apramycin and confirmed by PCR and sequencing. The recombinant R702 plasmid is then transferred into a suitable *E. coli* host by conjugation or electroporation, to create a methylating conjugal donor strain.

### Recombineering of Methyltransferase Genes of Interest

Along with the genome sequence, MTase genes of interest are identified using base modification data obtained from appropriate sequencing technologies, PacBio SMRT sequencing or Oxford Nanopore, and by accessing the Restriction Enzyme Database^27^. In the context of DNA transfer, a MTase gene of interest is such where a cognate and complete restriction endonuclease (REase) gene is present as a potential barrier to gene transfer.

The MTase gene is amplified along with 25 bp of the 5’ untranslated region (UTR), encompassing the putative Shine-Dalgarno (SD) ribosome binding site. For Type II and III systems, the modifying MTase gene component of the system is amplified, and for Type I RM systems, the modification (*hsdM*) and specificity (*hsdS*) genes are amplified together as one fragment, along with the native SD sequences. These genes are assembled into the insert region of a linearised pRBV1 vector, which, after sequence confirmation is amplified by PCR to result in a recombineering cassette PCR product.

The recombineering host *E. coli* strain “RE-InterStellar” was created by transference of the R702 plasmid and pKD46-Cm, a *catP* harbouring variant of pKD46 recombineering plasmid, into the *E. coli* “Stellar” *dam*^−^/*dcm*^−^ *recA*^+^ strain. The RE-InterStellar cells are primed for recombineering by induction of the lambda Red recombinase genes during competent cell preparation.

Amplified recombineering cassette linear DNA is transformed by electroporation into primed RE-InterStellar cells, which are recovered and plated on to LB medium supplemented with kanamycin and apramycin to positively select for successful integration of the recombineering cassette into the R702 plasmid. Integration at the *aadA1* locus of R702 results in the concomitant sensitivity to spectinomycin, which may be used as a negative screen. Following culture at 37 °C to lose the temperature sensitive pKD46-Cm plasmid, a confirmatory junction PCR, with primers spanning the site of integration, is performed (S 2). The recombinant R702 plasmid may now be mobilised, either be conjugation or electroporation, into an appropriate *E. coli* strain for subsequent transformation experiments (Figure 1).To ascertain the appropriate levels of arabinose induction the RFP reporter gene was inserted downstream of the P_BAD_ promoter and integrated into R702 and assayed for fluorescence (S 1).

**Table 1:** Bacterial Strains and Plasmids.

| Strains | Function/Characteristics | Source |
| --- | --- | --- |
| *E. coli* RE-InterStellar | Recombineering strain<br>*dam*<sup>-</sup>, *dcm*<sup>-</sup>, *recA*<sup>+</sup> | This study |
| *E. coli* sExpress | Conjugal strain<br>*dam*<sup>+</sup>, *dcm*<sup>-</sup>, *recA*<sup>+</sup> | Woods et al., 2019 |
| *E. coli* CA434 | Conjugal strain<br>*dam*<sup>+</sup>, *dcm*<sup>+</sup> | Mike Young |
| *E. coli* TOP10 | Cloning strain<br>*dam*<sup>+</sup>, *dcm*<sup>+</sup> | Invitrogen |
| *C. acetobutylicum* ATCC 824 | Solventogenic strain<br>Type II RM | ATCC |
| *C. difficile* R20291 | Hypervirulent outbreak strain<br>Type I RM, Type IV RM | Brendan Wren |
| **Plasmids** |  |  |
| pRBV1 | Recombineering base vector<br>Apr<sup>R</sup>, Em<sup>R</sup> | This study |
| pRBV1-Φ3TI | Recombineering base vector with inserted Φ3TI<br>MTase<br>Apr<sup>R</sup>, Em<sup>R</sup> | This study |
| pRBV1-HsdSM | Recombineering base vector with inserted<br>Cdi20291ORF16210P *hsdS* and *hsdM*<br>Apr<sup>R</sup>, Em<sup>R</sup> | This study |
| R702 | R Factor conjugal plasmid<br>Spec<sup>R</sup>, Tet<sup>R</sup>, Kan<sup>R</sup> | Hedges and Jacob, 1974 |
| R702-Φ3TI | R Factor plasmid with inserted Φ3TI MTase<br>Apr<sup>R</sup>, Tet<sup>R</sup>, Kan<sup>R</sup> | This study |
| R702-HsdSM | R Factor plasmid with inserted *hsdS/M* of<br>Cdi20291ORF16210P<br>Apr<sup>R</sup>, Tet<sup>R</sup>, Kan<sup>R</sup> | This study |
| pKD46-Cm | Lambda Red plasmid<br>Cm<sup>R</sup> | This study |
| pMTL83151 | Cm/Tm<sup>R</sup>, pCB102 Gram-positive replicon | SBRC |
| pMTL84151 | Cm/Tm<sup>R</sup>, pCD6 Gram-positive replicon | SBRC |

Spec<sup>R</sup> spectinomycin, Apr<sup>R</sup> apramycin, Em<sup>R</sup> erythromycin, Tet<sup>R</sup> tetracycline, Cm<sup>R</sup> chloramphenicol, Tm<sup>R</sup> thiamphenicol

### Transformation of *C. acetobutylicum* ATCC 824

The established method for transforming *C. acetobutylicum* ATCC 824 utilises the bacteriophage derived MTase, Ф3TI, on pAN1 and pAN2 plasmids to protect against restriction from the CAC824I Type II RM system (Figure 2 B)^28, 29^. To prove the concept of the presented methodology, Ф3TI was amplified from the pAN2 plasmid and inserted into R702 by the methods detailed, to yield R702-Ф3TI. The plasmid pMTL83151 was transformed into recombinant InterStellar cells harbouring R702-Ф3TI and subsequently extracted from induced cultures to be transformed into *C. acetobutylicum* ATCC 824 by electroporation. This was performed in parallel to the established method, transforming plasmids extracted from an *E. coli* TOP10 strain harbouring pAN2 and a negative no MTase control recombinant R702, R702-RBV1. Transformation efficiency is determined as a function of colony forming units (CFU) observed on CGM plates supplemented with thiamphenicol, per microgram of plasmid DNA. Transformation of plasmids extracted from the recombinant R702-Ф3TI strain was > 30 fold more efficient than those extracted from the R702-RBV1 harbouring strain (P < 0.0001) and at a statistically comparable transformation efficiency to the established method with TOP10 harbouring the pAN2 plasmid (Figure 2 A).

**Figure 2.**
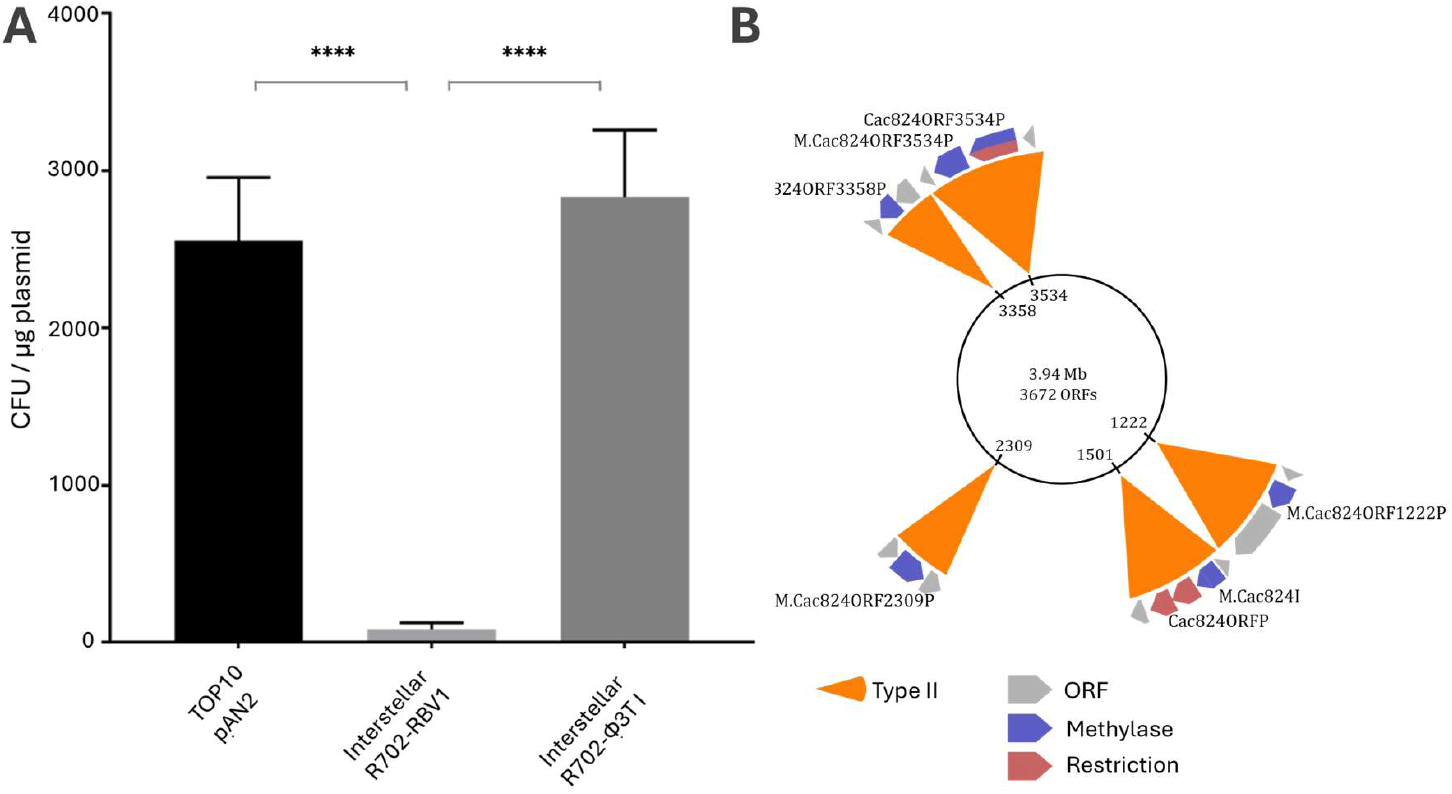
A: Transformation Efficiency of Plasmids Extracted from Different *E. coli* Hosts into *C. acetobutylicum* ATCC 824. All *E. coli* strains were cultured in parallel, in LB media supplemented with chloramphenicol and 1 mM L-arabinose for the strains harbouring recombinant R702 plasmids Interstellar R702-RBV1 and R702-pAN2 to induce expression of the heterologous Ф3T I MTase gene, driven by the AraC-P_BAD_ inducible system. Data obtained from TOP10-pAN2 extracted plasmids represents triplicate transformations, while data for the recombinant R702 harbouring InterStellar strains represents the average of four separate transformation experiments. Error bars represent the standard error of the mean (SEM) for each sample. **** represents statistical significance of P < 0.0001 between data sets. **B: The Restriction Modification Systems of *C. acetobutylicum* ATCC 824** The strain harbours five Type II RM systems, with one system of significance to DNA transfer, Cac824I, containing REase components which recognise and cleave DNA at the 5’-G<u>C</u>NGC-3’ sequence motif (adapted from The Restriction Enzyme Database, REBASE).

### Transformation of *C. difficile* R20291

The methodology was applied to the clinically relevant *C. difficile* ribotype 027 strain, R20291, of the lineage most closely related to the outbreak strain^30^. Consulting REBASE, two RM systems of relevance to DNA transfer were identified in the genome of R20291, a Type I and Type IV system (Figure 3 B). To overcome these barriers, the modification, *hsdM*, and specificity, *hsdS*, genes of the Type I system were amplified as a single PCR fragment and inserted, into R702, to yield R702-HsdSM. This recombinant plasmid was then transferred into *E. coli* Express (NEB), a strain lacking the *dcm* cytosine MTase gene to avoid host Type IV cleavage, yielding a methylating donor strain derivative of the previously described *E. coli* sExpress strain^24^. Conjugation of the plasmid pMTL83151 was performed in triplicate, using CA434 and two separate recombinant sExpress strains harbouring the control R702-RBV1 and R702-HsdSM, harbouring the modification and specificity components of the host Type I systems. Conjugation efficiency was measured as a function of CFU observed on BHIS plates supplemented with thiamphenicol, D-cycloserine and cefoxitin and total recipient CFU observed on BHIS plates without thiamphenicol selection. Conjugation efficiency of the R702-HsdSM harbouring sExpress donor strain was > 400 times higher than either of the control donor strains with P values < 0.001 (Figure 3 A).

**Figure 3.**
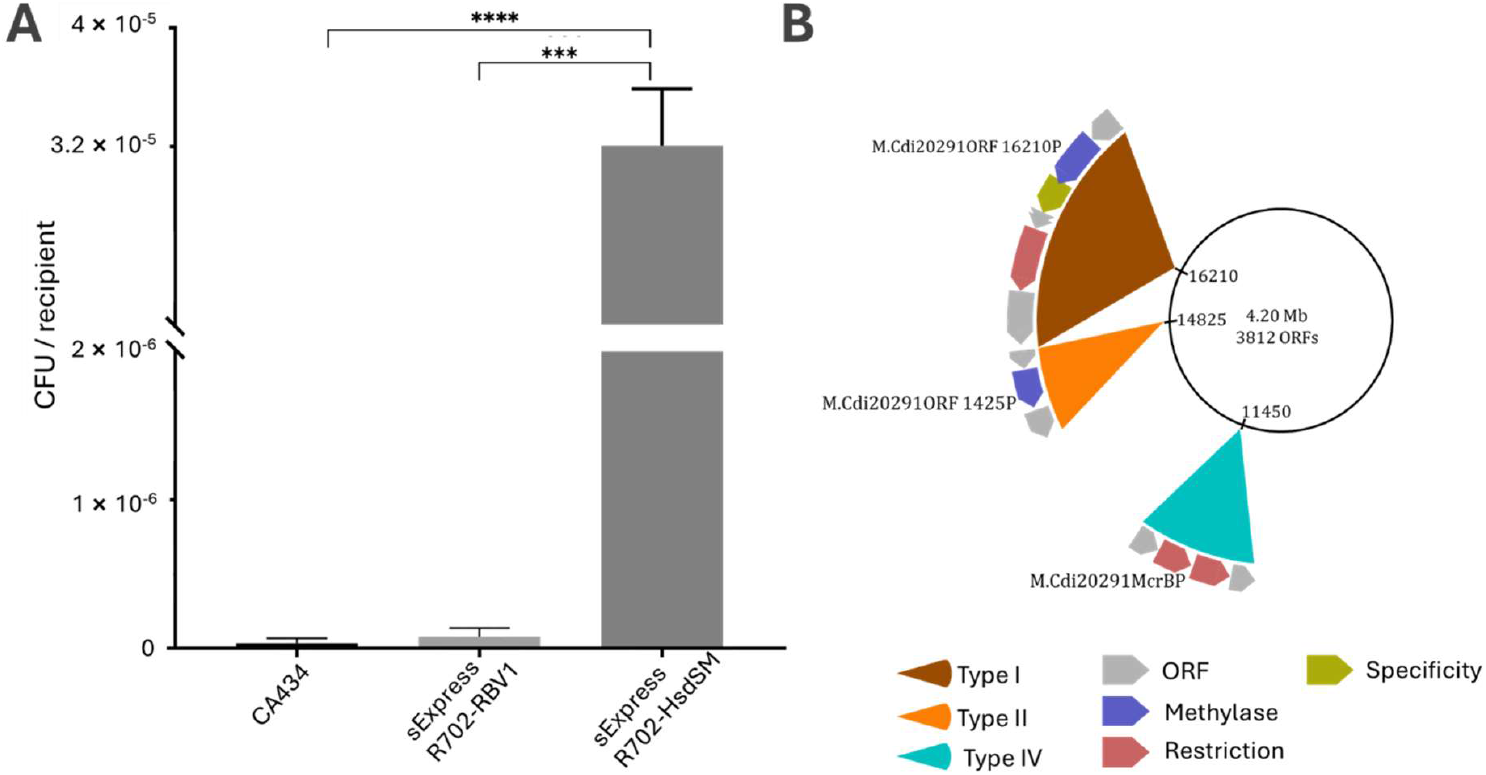
A: Conjugation Efficiency of *E. coli* Donor Strains into *C. difficile* R20291. The sExpress strain harbouring R702-HsdSM exhibited significantly higher conjugation efficiency than CA434 and the recombinant control sExpress strain harbouring R702-RBV1. The data plotted is representative of the average number of transconjugant colony forming units per recipient cell from biological triplicate experiments. Error bars represent the standard deviation of triplicate values. *** and **** represents statistical significance of P < 0.001 and P < 0.0001, respectively between indicated data sets. **B: The Restriction Modification Systems of *C. difficile* R20291** The strain contains one Type I, one Type II and one Type IV RM system. Of significance to DNA transfer are the Type I and Type IV systems (adapted from The Restriction Enzyme Database, REBASE).

## 3. Discussion

Genetic recalcitrance is a significant obstacle to the study of industrially and clinically important organisms. The strategy of harnessing methylation to protect exogenous transforming DNA has been utilised in a number of contexts to facilitate DNA transfer^21, 22, 23^. The novel methylating conjugal donor strains in this study were constructed by integration of protective methylation cassettes into the R702 conjugal plasmid at a defined locus, followed by transfer of recombinant R702 into a suitable *E. coli* host (Figure 1).

In the presented methodology, pertinent RM systems are identified in the genome of the organism of study and assigned sequence recognition motifs obtained through base modification data and utilising the REASE database. The modifying component of these systems is then inserted into the conjugal accessory plasmid, R702, utilising the lambda Red recombinase system via a recombineering base vector intermediate, pRBV1. The recombinant R702 plasmid is then transferred to an appropriate *E. coli* strain to create a methylating conjugal donor strain. Transfer of plasmid DNA can then be performed via conjugation or by electroporation of plasmids extracted from the recombinant methylating strains.

Considered choice of the appropriate *E. coli* donor strain genotype is important, particularly relevant in the context of the *dam*/*dcm* methylation status of *E. coli* strains, the presence or absence of which may be important to the evasion of host RM systems. This is dependent upon target host Type IV RM systems in the case of Dcm, or Type II RM systems in the case of Dam. The *E. coli* Dcm MTase methylates the internal cytosine of the recognition sequence 5’-C<u>C</u>WGG-3’, which is recognised and cleaved by Type IV Mcr cytosine-specific RM systems. The Dam MTase of *E. coli* modifies the internal adenine of the 5’-G<u>A</u>TC-3’ sequence motif, which may provide protection against certain Type II REases with overlapping recognition sequences to this motif, negating the requirement for a recombinant host MTase in that context. Both of these systems have been shown to affect DNA transfer from the *E. coli* hosts harbouring them^31^.

*E. coli s*train choice is also important for the recombineering step in the workflow. The Stellar strain, was identified as the most suitable host for recombineering. As an HST04 derivative, with a *dam*^−^/*dcm*^−^ genotype. The lack of the native Dam MTase, as well as the presence of a functional *recA* gene have been hypothesised to enhance the efficiency of lambda Red based recombineering^32, 33^. Harbouring both the pKD46-Cm recombineering plasmid and R702 conjugal plasmid, the strain was designated, “RE-InterStellar”.

The efficacy of the methodology was assessed in the transformation of the model solventogenic organism, *C. acetobutylicum* ATCC 824. This proof of concept was based upon the established method, using the *B. subtilis* bacteriophage derived MTase Ф3TI on multicopy plasmids, to enable DNA transfer into this industrially relevant organism^28, 29^. The cytosine methyltransferase, Ф3TI, modifies the internal cytosine of the recognition sequences 5’-GG<u>C</u>C-3’ and 5’-G<u>C</u>NGC-3’^34^. The internal 5-methylcytosine in the latter recognition sequence results in protection from cleavage by the native Cac824I REase (Figure 2B). There are 22 instances of the GCNGC sequence motif present in the shuttle vector pMTL83151, rendering this plasmid eminently vulnerable to restriction by the CAC824I system. This methodology exhibited transformation efficiencies comparable to the established method, proving the efficacy of induced P_*BAD*_ directed expression of a MTase gene from the single copy R702 plasmid.

Next the methodology was exemplified for the clinically important *C. difficile* strain, R20291 outbreak strain. Classified as belonging to the BI/NAP1/ PCR ribotype 027 groups and designated as ‘hypervirulent’ due to the ability of this strain to produce higher titres of toxin, an additional binary toxin and elevated numbers of spores^35^. This strain harbours both Type I and Type IV RM systems (Figure 3B), based upon the REBASE entry for this strain and 100% homology of the Cdi20291ORF16210P system to a closely related 027 strain for which PacBio base modification data has been obtained (data not shown), the targeting motif 5’-C<u>A</u>NNNNNNN<u>T</u>AAAG-3’ is assigned to the Type I system. There is a single binding motif for this system in the pMTL84151 plasmid and as such a synergistic effect between this system and the McrBC family Type IV system is likely to be impeding DNA transfer in this strain. Use of the recombinant R702 in this context, and the *dcm*^−^ *E. coli* donor strain sExpress, resulted in greater than 400 times higher conjugation efficiencies over the control strains. These data further validate the presented method and also provides a mechanistic insight into Type I RM systems, the hetero-oligomeric protein complex is able to perform methylation activity without the restriction component^35^.

It has previously been reported that the *C. difficile* R20291 strain used in this work, most closely related to clinically relevant outbreak strain, exhibits lower conjugation efficiencies than R20291 strains from other sources^29^. Variant analysis of the sequenced genomes revealed a number of single nucleotide polymorphisms (SNPs) in these other strains, resulting in significantly attenuative phenotypic differences in terms of key virulence factors, but higher efficiency of DNA transfer. Whilst not directly associated with the RM systems highlighted in this study, it is hypothesised that the presence of functional peritrichous flagella in the closest ancestral strain could impede the conjugation process, further complicating DNA transfer in this strain. The method presented in this work therefore represents an appropriate way of studying the strain closest to the index case strain and of significant clinical importance.

This strategy enables an increase in efficiency transfer of DNA into bacteria, genetically recalcitrant due to the presence of host RM systems. Integration into R702 represents a more efficient approach to overcoming the restriction barrier by conjugation, advantageous by the ease at which recombinant R702 harbouring MTase genes may be transferred to a suitable *E. coli* strain, creating a methylating conjugal donor strain. The methylating conjugal donors created with this methodology avoid the need for resynthesis of plasmid components to silence restriction sites and the complications of tri-parental mating methods. Crucially, with this strategy, there are no requirements to vitiate the organism of study with RM system mutations or damaging methods such as heat shock, enabling study of the strain closest to the wildtype context, particularly important to clinically relevant organisms such as *C. difficile* and other pathogens.

## 4. Materials and Methods

### 4.1 Growth of Bacterial Strains

*E. coli* strains were cultured in LB broth and agar plates. Cultures were incubated aerobically at 37 °C with 200 RPM orbital shaking for liquid cultures. The *E. coli* recombineering host strain, RE-InterStellar, was incubated at 30 °C to maintain the temperature sensitive pKD46-Cm plasmid. *Clostridium* and *Clostridioides* strains were incubated anaerobically in a Don Whitley workstation with an atmospheric composition of 80% nitrogen, 10% carbon dioxide, 10% hydrogen at 37 °C. *C. acetobutylicum* ATCC 824 was cultured in 2 × YTG broth (pH adjusted to 5.2 prior to autoclaving) and on CGM agar plates (pH adjusted to 7 prior to autoclaving). *C. difficile* R20291 was cultured in BHIS broth and agar plates, BHI supplemented with 5 g L^−1^ yeast extract and 1 g L^−1^ L-cysteine.

Growth media were supplemented with antibiotics, where appropriate at the following concentrations: *E. coli*: chloramphenicol (25 μg mL^−1^), spectinomycin (100 μg mL^−1^), kanamycin (50 μg mL^−1^) apramycin (100 μg mL^−1^); *C. acetobutylicum* and *C. difficile*: thiamphenicol (15 μg mL^−1^) and D-cycloserine (250 μg mL^−1^), cefoxitin (8 μg mL^−1^) to select against *E. coli* donor cells.

### 4.2 *C. acetobutylicum* Transformation

Plasmids to be transformed were extracted from recombinant R702 harbouring *E. coli* overnight culture with 1% v/v 1 M L-arabinose supplementation and the appropriate controls strains. Transformations were performed using a GenePulsar XcellTM electroporator, with both CE and PC modules (BioRad). Competent cell aliquots were added to 500 ng of plasmid DNA, and incubated for 2 minutes on ice. Samples were then electroporated using the following settings: exponential wave, voltage 2.0 kV, resistance ∞ Ω, capacitance 25 μF. Pulsed samples were transferred to 10 mL fresh 2 × YTG medium and recovered for 4 hours. Samples were then centrifuged at 5000 × *g* and plated onto selection plates with appropriate antibiotic selection.

### 4.3 *C. difficile* Conjugation

Donor *E. coli* strains harbouring recombinant R702 and the plasmid to be transferred, were set up in parallel to overnight *C. difficile* cultures, with 1% v/v 1 M L-arabinose added to donors where required for expression of MTase genes, along with control strains. 1 mL of donor strain overnight cultures were centrifuged at 4000 × *g* for 2 minutes and supernatant removed, before washing with 500 μL PBS. Donor pellets were then transferred to the anaerobic cabinet. The pellets were resuspended in 200 μL of *C. difficile* overnight culture and spotted onto BHIS mating plates. Mating plates were left overnight before growth was harvested by flooding plates with 1 mL PBS, serially diluted and transferred to plates with appropriate antibiotic selection.

### 4.4 Lambda Red Primed Electro-Competent *E. coli* Cells

Electro-competent *E. coli* RE-InterStellar cells were prepared by established methods, except for incubation at 30 °C and antibiotic supplementation; chloramphenicol (25 μg mL^−1^), kanamycin (50 μg mL^−1^), to maintain pKD46-Cm and R702 plasmids respectively and the addition of 1% v/v 1 M L-arabinose at OD_600_ of 0.1 to induce expression of the lambda Red recombinase. Prepared electrocompetent cells were stored at −80 °C prior to use.

## Supplementary Information

### S 1 Arabinose Inducible System Promoter Assay

An RFP based promoter assay was performed to confirm the efficacy of the AraC-P_BAD_ promoter system in the R702 context and appropriate inducer concentration. The recombineered rIS-RFP strain was assayed with a range of L-arabinose induction. Fluorescence was measured at 15.5 hours, representative of expression following an overnight growth cycle, as would be the case for methylase expressing cassettes prior to plasmid extraction or conjugation. The strains were also assayed at 24 hour time points.

**Figure S1.**
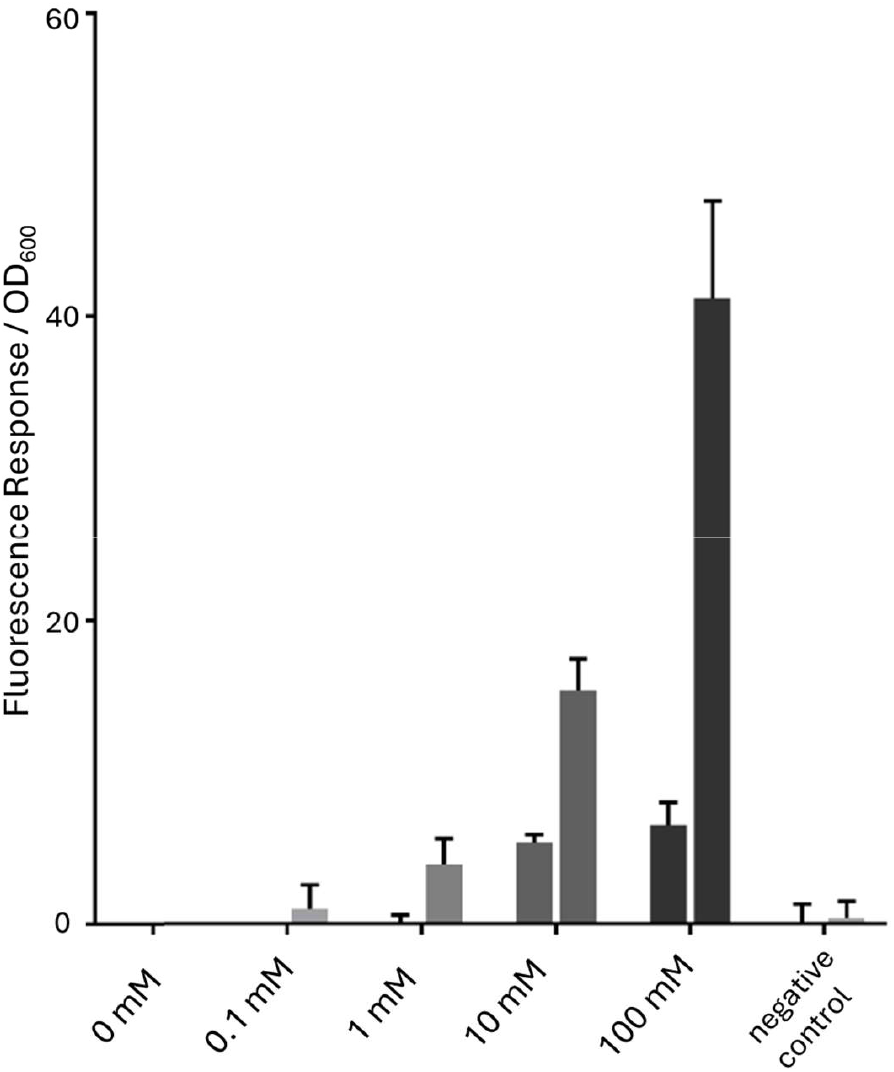
RFP Assay of Recombinant AraC-P_BAD_ Cassette. Re-IS-RFP samples cultured in a 96 well plate format with constant shaking at 30 °C with varying levels of L-arabinose induction. Bar pairs represent the average fluorescent response adjusted for OD_600_ for triplicate samples at 15.5 hour and 24 hour timepoints for each sample, represented by the first and second columns for each sample, respectively. The RBV control values are representative of six replicates. Error bars represent the standard deviation of fluorescent response from triplicate samples of induced rIS-RFP and the six replicates of the RBV1 control sample. Fluorescence was measured at a wavelength of 584 nm wavelength and an emission detection at 621 nm.

**Table S2.**
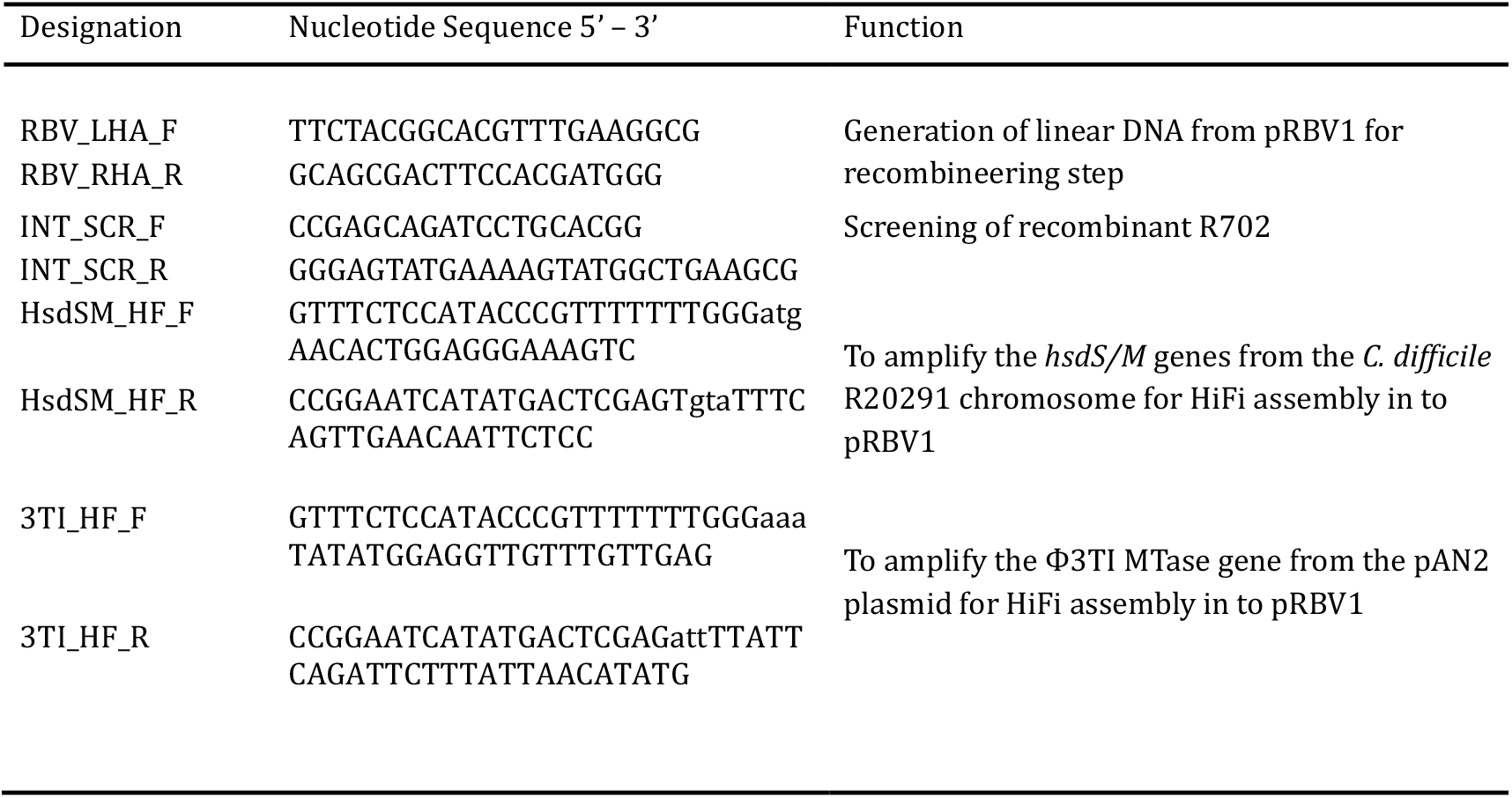
Primers.

